# Cryptic multicellularity in wild yeast brings selective advantages under stress

**DOI:** 10.1101/2025.11.06.687064

**Authors:** W. K. Sexton, K Schmidt, Q Dickinson, J Childress, F Rosenzweig, E Kroll

## Abstract

Multicellularity has evolved dozens of times in many branches across the Tree of Life, in each case producing a new kind of individual that has the potential for division of labor among its constituent cells. In many of these instances, nascent multicellularity has remained facultative, manifesting only under conditions where it provides a decisive fitness advantage to the genome in which it has arisen. Here we investigate the mechanistic basis for, and the selective advantages of, cryptic multicellularity in wild strains of the yeast *Saccharomyces cerevisiae.* Meiotic purification of this trait, followed by genome sequencing, indicates that the genome of diploid champagne yeast DB146 is homozygous at the *AMN1* locus (<u>A</u>ntagonist of <u>M</u>itotic exit <u>N</u>etwork) for an allele that dysregulates post-mitotic cell separation. Expression of this trait in haploid derivatives of DB146 results in clonal multicellular clusters in which daughters remain attached to mother cell walls. Systematic analysis of viability in haploid and diploid DB146 derivatives subjected to benign conditions as well as to starvation, desiccation, and low temperature at different cell titers demonstrates that haploid multicellular variants exhibit higher survivorship under conditions that wild yeast likely experience when they overwinter. Examining a collection of other wild strains, we find that ∼20% exhibit ploidy-dependent expression of clonal multicellularity. Altogether, our data suggest that in yeast that sporulate when starved, cryptic multicellularity may come under balancing selection because haploid multicellular progeny enjoy a transient survival advantage that fades once favorable growth conditions resume.

## Introduction

The study of multicellularity (MC) in the budding yeast *Saccharomyces cerevisiae* has opened a window on one of life’s major innovations, the transition from single cells to multicellularity (1,2). Yeast can achieve multicellularity either by unicells coming together (aggregative multicellularity) or staying together (clonal multicellularity) (3). Aggregative multicellularity in yeast is triggered by changes in adhesive properties at the cell surface, leading to flocculation (4), whereas the emergence of clonal multicellularity initially stems from a defect in mother-daughter separation, leading to the formation of cell clusters (5,6). If novel clonal multicellular variants are to compete successfully with their single-celled ancestors, they must initially bear traits that enhance their survival and/or reproductive capacity (7,8). After all, developing heritable advantages associated with division of labor would almost certainly require many additional rounds of mutation and selection (9). Among the traits that one might expect in any nascent multicellular organism is enhanced stress resistance due to physical protection offered by diffusion barriers or physiological heterogeneity within an otherwise clonal body. Among facultatively multicellular organisms like the slime mold, *Dictyostelium discoideum*, stress in the form of nutrient limitation serves as a signal that triggers aggregation and differentiation (10) – though aggregation in this species often produces a chimera rather than a clone (11). Diverse forms of facultative multicellularity exist across the Tree of Life among organisms most often viewed as unicellular (12). In species having the capacity to become multicellular as part of their life cycle, the decision to do so is usually triggered by some type of stress (13).

Heritable clonal multicellularity can emerge under artificial selection in the laboratory. For example, in *Saccharomyces cerevisiae*, multicellular clusters readily evolve under centrifugation (settling) selection, demonstrating that this evolutionary transition can occur within a few score generations in the lab under the right conditions, however unnatural (1,5). Under further settling selection, these clusters begin to exhibit simple life cycles and even a primitive division of labor, where some cells undergo apoptosis, apparently for the benefit of the whole (14). Other, more ecologically realistic forms of selection can also favor the evolution of multicellularity from single-celled ancestors. Predation, which selects for large prey size, can trigger colony formation in the single-celled green alga, *Chlorella vulgaris* (15), and can favor the evolution of stably heritable multicellularity in experimental populations of *Chlamydomonas reinhardtii* (16). Also, spatially limited resources have been theoretically shown to favor the emergence of multicellularity at a critical resource size (17). Thus, rudimentary multicellular bodies offer immediate ecological advantages through enhanced survival and metabolic efficiency (18), potentially providing a competitive edge in environments with predictable sources of mortality and/or unpredictable availability of limiting resources.

Here, we demonstrate in yeast how different forms of stress can facilitate the transition from single cells to clonal multicellularity by showing that a multicellular lifestyle can be advantageous under diverse ecologically relevant conditions. The type of multicellularity described here is cryptic, as its expression is ploidy-dependent, and it is based on mutations in *AMN1,* which governs post-mitotic mother-daughter cell separation (19,20). Additional support for our findings emerges from surveys (presented here and in (21)) showing that many strains of wild yeast harbor cryptic multicellular phenotypes that are expressed in a ploidy-dependent manner. The evolution of multicellularity, therefore, depends on both the contents of a lineage’s genetic toolkit and the environment that that lineage experiences, highlighting the need for a deeper – and broader – understanding of the various mechanisms by which multicellular life can arise and be perpetuated.

## Methods

### Strains, genetic manipulations and growth conditions

The starting point for the experiments described here is yeast strain DB146 (a gift from B. Blondin and F. Vezinhet). DB146 is a mitochondrially marked derivative of diploid strain 8010, which was isolated from a champagne vineyard (22). DB146 exhibits chromosomal length polymorphism (22,23), the dynamics of which is markedly influenced by starvation (24). All strains used in this work and their provenance are listed in Supplementary Materials, Table S1. Yeast cultures were incubated on solid and liquid YEPD (1% glucose) media at room temperature (24°C). Meiosis was induced by plating cells on yeast sporulation medium (1.0% potassium acetate, 0.1% yeast extract, 0.05% glucose, 2.0% agar), followed by incubation at room temperature for 5-7 days (25). Tetrad dissection was performed using a SporePlay dissection microscope (Singer Instruments, Roadwater, Watchet TA23 0RE, United Kingdom). Allele replacement was performed essentially as in (26).

Ploidy and mating type of yeast strains were determined via colony PCR. The primers used were 5’-AGTCACATCAAGATCGTTTATGG-3’ (“MAT1”), which targets outside of the MAT locus, 5’-GCACGGAATATGGGACTACTTCG-3’ (“MAT2”), which is specific for MATα, and 5’-ACTCCACTTCAAGTAAGAGTTTG-3’ (“MAT3”), which is specific for MATa (27). The reaction mixture consisted of 1x DreamTaq PCR Master Mix (Thermo Fisher Scientific, Waltham, MA, USA), 1 µM of each primer, and 0.2 µg/mL BSA. Yeast colony matter suspended in sterile deionized water and heated at 95 °C for 15 min was used as a template. The PCR temperature profile consisted of an initial step of 95 °C for 3 min, 35 cycles of 95 °C for 30 sec, 58 °C for 30 sec, and 72 °C for 1 min, and a final elongation step of 72 °C for 5 min. PCR products were visualized via gel electrophoresis using 1% agarose gel in 1x TAE buffer with 0.5 µg mL^-1^ ethidium bromide.

### Microscopy

Unicellular and multicellular yeast strains were imaged by light and by epifluorescence microscopy using a Zeiss Axioskop microscope, as well as by transmission electron microscopy (TEM) and scanning electron microscopy (SEM). For TEM, cells were fixed for 1 h in 3% glutaraldehyde in cacodylate buffer, washed once in cacodylate buffer, resuspended in 1% osmium tetroxide and 1% potassium ferricyanide in cacodylate buffer, and incubated for 30 min at 23 °C. Cells were then washed several times in dH2O, resuspended in 1% thiocarbohydrazide in water, and incubated for 5 min at 23 °C. Cells were again washed in dH2O, incubated in 1 % osmium tetroxide and 1% potassium ferricyanide in cacodylate buffer for an additional 5 min, and washed again in dH2O. Treated cells were then incubated in saturated uranyl acetate for 2 h and dehydrated through a graded series of acetone washes. The dehydrated samples were embedded in Epon 812, sectioned, and imaged at 80 kV.

For SEM, cells were placed in 2% EM-grade glutaraldehyde in cacodylate buffer, pH 7.2, fixed overnight at 4°C, then rinsed with distilled water (dH_2_O) and placed in a solution of 2% osmium tetroxide (OsO4) in dH2O for 1 hour at 25 °C. OsO_4_ was rinsed off via several washes with cacodylate buffer, pH 7.2. Cells were then dehydrated via a succession of 10-minute incubations in 40%, 50%, 70%, 90%, and 95% ethanol, followed by two 10-minute washes in 100% ethanol. Ethanol was removed and replaced with 2 rinses of 100% propylene oxide (PO), whereafter samples were infiltrated with a 1:1 solution of PO and EmBed 812-DER 736 epoxy (Electron Microscopy Sciences) for 4 hours at 25 °C. This solution was removed and replaced with 100% EmBed 812-DER 736 and incubated overnight at 25 °C. Samples were then cured at 60 °C for 24 hours, whereupon the sample block was trimmed and sectioned using an RMC MTXL ultramicrotome. Sections collected on grids were stained with uranyl acetate and lead citrate, air dried and then imaged using either a Hitachi H-7100 transmission electron microscope at 75 kV or a Hitachi S-4700 Type II cold field emission scanning electron microscope, using procedures described by (28) (Hitachi Group, Schaumburg, IL, USA).

### Criteria for distinguishing multicellular from unicellular yeast strains

Multicellularity in yeast and other eukaryotic microbes can be aggregative or clonal (3). In *Saccharomyces*, the mechanistic basis for the former is flocculation, which can result in a genetically heterogeneous multicellular aggregate. In contrast, the basis for the latter is typically the failure of mother and daughter cells to separate, resulting in a genetically homogenous “organism,” albeit one that may vary orders of magnitude in cell number and size (2). Two assays were used to distinguish multicellular from unicellular yeast: (1) treating cell clusters with EDTA, which breaks up genetically heterogeneous cell aggregates (“flocs”), treating cell clusters with zymolyase, which dissolves cell wall bridges of cells that fail to separate following cytokinesis (29). Clusters that were not dissociated by EDTA, but that were dissociated by zymolyase, were deemed to be truly multicellular. Multicellular and unicellular yeast in our experiments could also be distinguished by colony morphology: multicellular colonies appeared rough, while unicellular colonies appeared smooth (see **Supplementary Materials, Figure S1**).

### Analysis of cell and cell cluster size

The size distributions of yeast cells and cell clusters were measured using a Multisizer 4e particle analyzer (Beckman Coulter, Brea, CA, USA). Isolates were grown overnight in liquid YPD and then diluted 1000x in ISOTON II (Beckman Coulter, Brea, CA, USA). 0.5 mL of the dilutions was sampled using the 280 µm aperture.

To characterize the size distributions of unicellular (UC) and multicellular (MC) derivatives from DB146, seven technical replicates were used per isolate, with roughly 3 × 10^4^ to 7 × 10^4^ bodies (cells or cell clusters) sampled per replicate. For each size bin, the differential volume from all seven replicates was averaged. To test the ability of the particle analyzer to resolve different size distributions, mixtures were prepared with varying proportions of a UC (5) and MC (7) derivative from DB146. Cell and cell cluster titers of overnight YPD cultures were determined using a hemocytometer and light microscopy. Suspensions with different proportions of the MC isolate were prepared (0%, 5%, 10%, 15%, 20%, 40%, 45%, 50%, 55%, and 60%) to a uniform concentration of 3 × 10^7^ cells and/or clusters per mL. Aliquots of these cell suspensions were diluted in ISOTON II as described above and run in triplicate. The size of 5010 bodies (cells or clusters) for each replicate was measured and used to calculate weighted medians (S50) for each mixed culture. Bootstrap resampling was done with each dataset to obtain 95% confidence intervals for each calculated weighted median.

### Assessment of growth rate and yield under nutrient-replete conditions

Growth rate and yield were estimated by measuring optical density at 600 nm (OD_600_) using a BioTek Synergy HXT plate reader (Agilent Technologies, Winooski, VT, USA). Yeast populations were grown in YPD, 1% glucose at 25 °C in a 48-well microtiter plate format (500 µL). Mean values for each parameter were computed from a minimum of three independent replicates. Demographic parameters consisted of maximum specific growth rate (µ_max),_ derived from the slope of log-phase growth (where r^2^ =∼0.995 or greater) and yield at stationary phase (Y_s_), estimated as OD_600_ after 48 hours.

### Assessment of survivorship under starvation conditions

To determine survivorship under nutrient-deficient (starvation) conditions, triplicate populations were established in YPD (1% glucose), and the fraction of viable cells remaining was estimated at successive timepoints over the course of 100 days. Yeast cells were incubated without shaking on the benchtop at room temperature in sealed 48-well plates. Cells in starving populations were assayed for viability every 5-10 days using Trypan blue exclusion, as described by (30).

### Estimate of relative fitness under multiple ecologically relevant conditions

Relative fitness of unicellular and multicellular yeast was estimated under conditions that differed with respect to: (1) NUTRIENT CONTENT (YPD 1% glucose or distilled H_2_O), (2) TEMPERATURE (24 °C or 10 °C), (3) INITIAL DENSITY (5 × 10^7^ cells mL^-1^ or 1 × 10^4^ cells mL^-1^), (4) CULTURE MODE (continuously hydrated or periodically desiccated), (5) INITIAL PROPORTION OF MULTICELLULAR AND UNICELLULAR STRAINS (9:1 MC:UC or 1:9 MC:UC). Pure cultures of each of several independent MC isolates (44.26, 46.2, 46.16 and MC7) and the UC ancestor (DB146) were pre-grown separately for 24 hours at 24 °C under nutrient-replete conditions (YPD, 1% glucose) In all but condition (5), MC and UC strains were mixed at time-zero in a 1:1 ratio and slowly shaken to prevent sedimentation. Each MC:UC mixed culture and control “pure” MC and UC cultures were set up as a 5 mL culture in a 15 × 100 mm glass tube with a slide cap; mixed cultures were set up in triplicate, pure cultures in duplicate. Cultures were slowly agitated to prevent sedimentation. Cultures subject to periodic desiccation (4) were incubated for one day in 500 µL liquid YPD medium without shaking. The next day, after all cells had settled, 400 µL of supernatant was removed, leaving a suspension of ∼100 µL medium + cells, which, after drying out, was kept desiccated for 15 days. Desiccated cultures were rehydrated with 500 µL dH_2_O, and a sample was plated to determine the MC:UC ratio. A 400 µL supernatant of this cell suspension was again removed to let the cell culture desiccate until the next round of counting.

Under conditions 1, 2, 3, 4, and 5 and their combinations, relative fitness was deduced from the change in the ratio of multicellular evolved strains to their unicellular ancestor (MC:UC) over 45 days of starvation in three intervals of 15 days each. To estimate the change in MC:UC ratios, aliquots of the experimental populations were plated on YPD agar at days 0, 15, 30 and 45. Following incubation for 3 days at 24°C, the number of colonies formed on these plates was counted and their respective morphologies (smooth=UC, wrinkled=MC) recorded. Duplicate unmixed MC and UC “pure culture” controls were processed identically to compare the viability of pure and mixed cultures. Proportions of UC and MC colonies were converted to selection rate constants per day using the formula 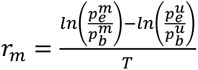, where 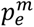 is the proportion of MC colonies at the end of the time period T (15, 30, or 45 days), 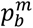 is the proportion of MC colonies at the beginning of the period T (0, 15, 30 days, respectively), and 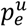 and 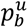 are the same for UC colonies. Overall, 121,095 colonies were analyzed across all experiments.

### Meiotic purification and genomic sequencing

A representative haploid (mating type <u>a</u>) clone of an MC strain (MC7) was crossed to the haploid (mating type alpha) reference UC strain BY4742 (https://www.yeastgenome.org/strain/by4742), and multiple diploid colonies were confirmed by mating-type locus PCR (27). Following sporulation of these MC7 x BY4742 diploids, a multicellular MAT-a haploid derivative (F1) was chosen and crossed again to BY4742. Selection of multicellular MAT-a haploids and mating to the unicellular reference strain was repeated 6 times, effectively using meiosis and backcrossing to isolate genetic determinant(s) of the multicellular phenotype. Four F6 progeny were sequenced via Illumina NextSeq and compared to the genome of parental strain BY4742 as follows: the genome sequences of BY4742 and DB146 were concatenated to create a single file. The four backcrossed isolates were sequenced using Illumina MiniSeq at ∼20X coverage. Illumina reads were trimmed with Trimmomatic and aligned to the concatenated genome using Bowtie2. Reads were then filtered with Samtools based on their MAPQ score. All aligned reads having an MAPQ score < 4 were discarded, whereafter coverage of the filtered alignment was determined using deepTools (31). All listed analyses were performed on the public Galaxy web server at usegalaxy.org (32). The Integrative Genomics Viewer (33) was used to compare the coverage maps to the BAM files and to determine which DB146 regions had been introgressed into the BY4742 genome to produce the multicellular phenotype.

## Results

### Sporulation uncovers a clustering phenotype in a hemithallic wild yeast strain

Random spore analysis of champagne yeast DB146 (22) showed that approximately half of its meiotic progeny formed cell clusters. Cell clusters ranged from 4 to >100 cells with cells fully divided but attached to each other (**Figure 1**). Tetrad dissection analysis confirmed that the ratio of unicellular (UC) to clustered, or multicellular (MC), meiotic progeny was 2:2, indicating that inheritance of the multicellular phenotype was consistent with an apparent Mendelian pattern of inheritance (*data not shown*).

**Figure 1.**
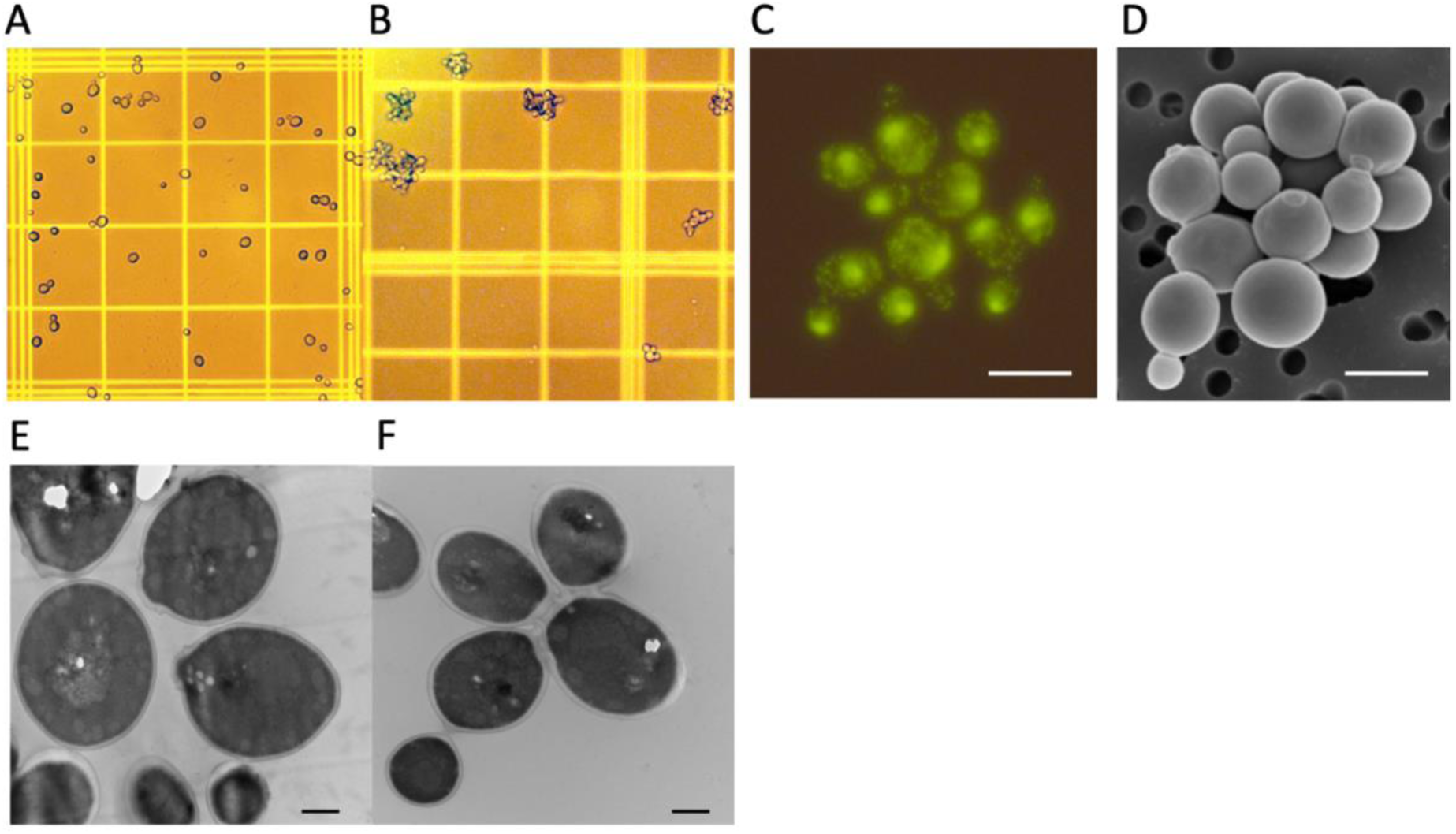
Clustering phenotype in meiotic derivatives of DB146. **A:** Single cells in a culture of ancestral strain, DB146; **B:** Clustering in a DB146 meiotic derivative. A and B are imaged on a standard hemocytometer. **C**: A cell cluster with its DNA stained using CyberGreen. **D:** A cell cluster viewed by SEM. For C and D, the bar is 5μ. **E**, **F**: Unicellular diploid ancestor (E) and multicellular haploid derivative (F) viewed by TEM. For E and F, the bar is 1μ.

Particle analysis of resulting cell size distributions exhibited broader distributions in volume than single-cell cultures with cluster size maxima exceeding those of single–celled cultures 2- to 3- fold (**Figure 2 A,B**). Upon checking the mating type of clustered and single-celled spore progeny of DB146, we found that half of DB146 offspring were clustered and haploid, while the other half were unicellular and diploid (**Figure 2C**). Since spore progeny diplodize only in the presence of a functional *HO* allele (35), the 2:2 segregation pattern of haploid and diploidized spore progeny suggested that one of two *HO* alleles in the diploid ancestor DB146 might be inactive, rendering DB146 a “hemithallic” strain, in analogy with homothallic yeast strains which contain two working *HO* alleles, and heterothallic strains, which contain two inactive *HO* alleles (34). Sequencing the *HO* locus confirmed that one of two *HO* alleles in the diploid ancestor DB146 contained a premature stop codon along with several other mutations, rendering this *HO* allele inactive (data not shown). Since the novel clustering phenotype co-segregated with ploidy, we assumed that the clustering, or multicellular phenotype, is either ploidy- or gene dosage-dependent and is caused by a homozygous mutation in the DB146 ancestor.

**Figure 2.**
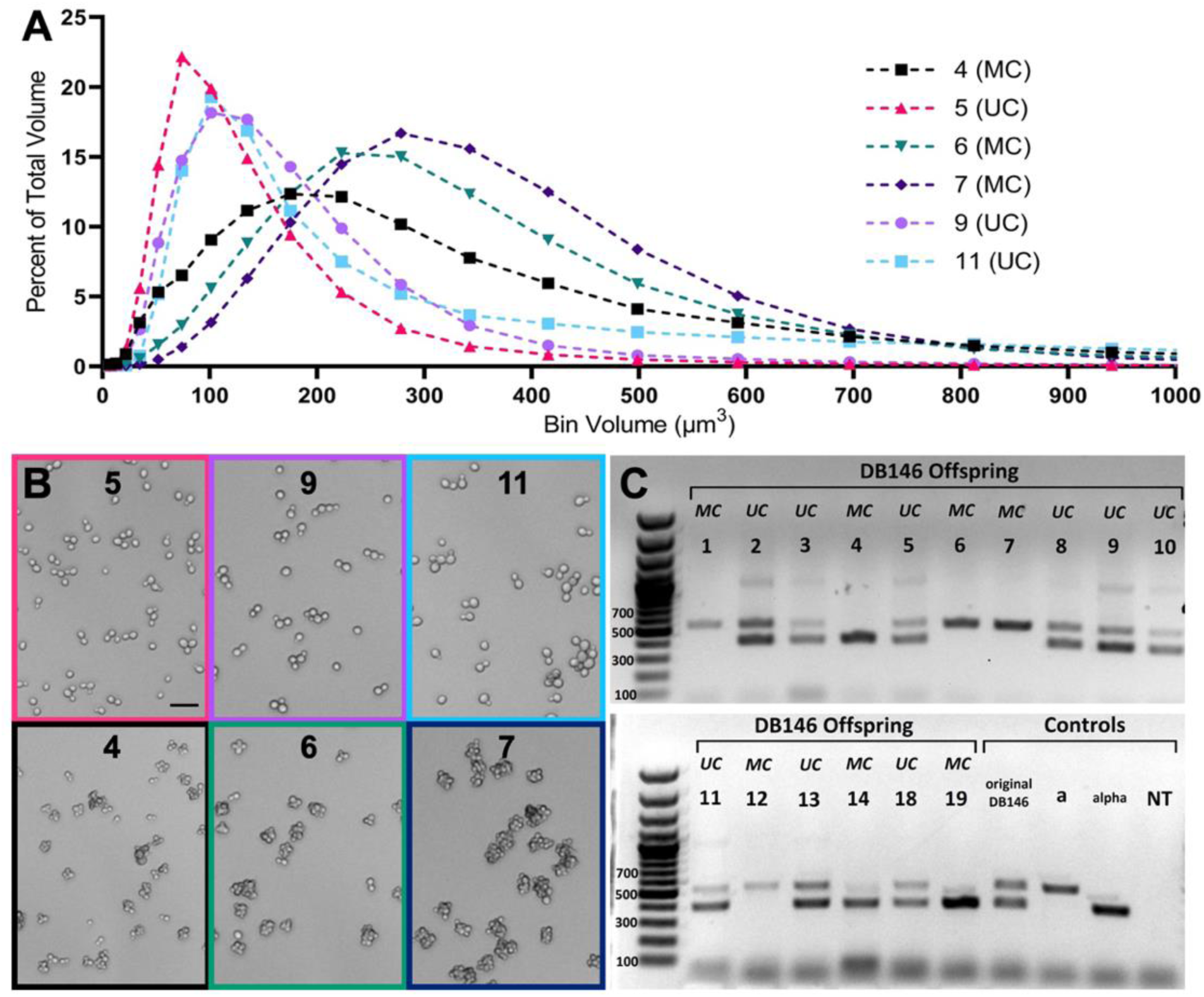
Demography, morphology and ploidy of unicellular and multicellular yeast cultures. **A:** Size distributions of three representative unicellular (UC) (5, 9, 11) and three multicellular (MC) (4, 6, 7) derivative populations. **B:** Photomicrographs of cells depicted in panel A; scale bar length = 20μm. **C:** Ploidy of individual isolates determined by PCR of single colony isolates targeting the *MAT* locus. Single band indicates haploidy, while double bands indicate diploidy. Irrespective of mating type, all diploid isolates are UC, while all haploid isolates are MC.

To determine whether cell clusters formed as aggregates by cell adhesion (i.e., by flocculation (35)) or as clones by failure of mother-daughter separation (36) clusters were first vigorously vortexed and then sonicated in the presence of EDTA, which helps to dissolve “flocs” by chelating Ca^2+^ bridges (37). Cell clusters remained intact following this procedure. By contrast, digestion with zymolyase, which digests fungal cell walls, converted cell clusters into single cells, indicating that clusters originate as clones.

To better understand the physical basis for clustering, the clustering phenotype was investigated by scanning electron microscopy (SEM) and transmission electron microscopy (TEM). SEM showed that cell walls of clusters were more resilient to sample preparation than those of the unicellular ancestor, which crumpled under the harsh conditions of sample preparation (**Fig. 2S**). This finding suggested that cell walls might be thicker in clustered cells. TEM confirmed this, as cells in clusters had thickened cell walls and robust intercellular bridges consistent with a defect in cell wall maintenance and cell separation following cell division (**Figure 1**: panels **E, F, G**, and **H**). To summarize, we concluded that cell clusters in multicellular haploid isolates of strain DB146 arise clonally from a single cell whose progeny fail to separate after cell division. Hereafter, we frequently refer to strains that propagate as clonal clusters as <u>m</u>ulti<u>c</u>ellular (MC), the single-celled ancestor lacking this phenotype, and all other single-celled strains as <u>u</u>ni<u>c</u>ellular (UC). Additionally, we conceive clonal, non-aggregative multicellular clusters as ‘bodies,’ rather than cells, and refer to them as such.

### The extent to which a culture is multicellular can be ascertained by S50, an index that represents the central value of a population’s biomass

Multicellular populations consist of a mix of different cluster sizes, with most of a population’s biomass concentrated in physically large but numerically infrequent clusters. To quantify the degree of multicellularity, we developed an index, the weighted median S50, representing the central value of a population’s biomass. S50 is the characteristic size (by volume) of a cell cluster at which the mass of all cells in clusters exceeding S50 comprises 50% of the total population volume. Our index is analogous to the sequencing quality parameter N50 (38). This descriptive statistic for cluster size distribution can be used to distinguish cell populations based on cluster size (**Figure 3**).

**Figure 3.**
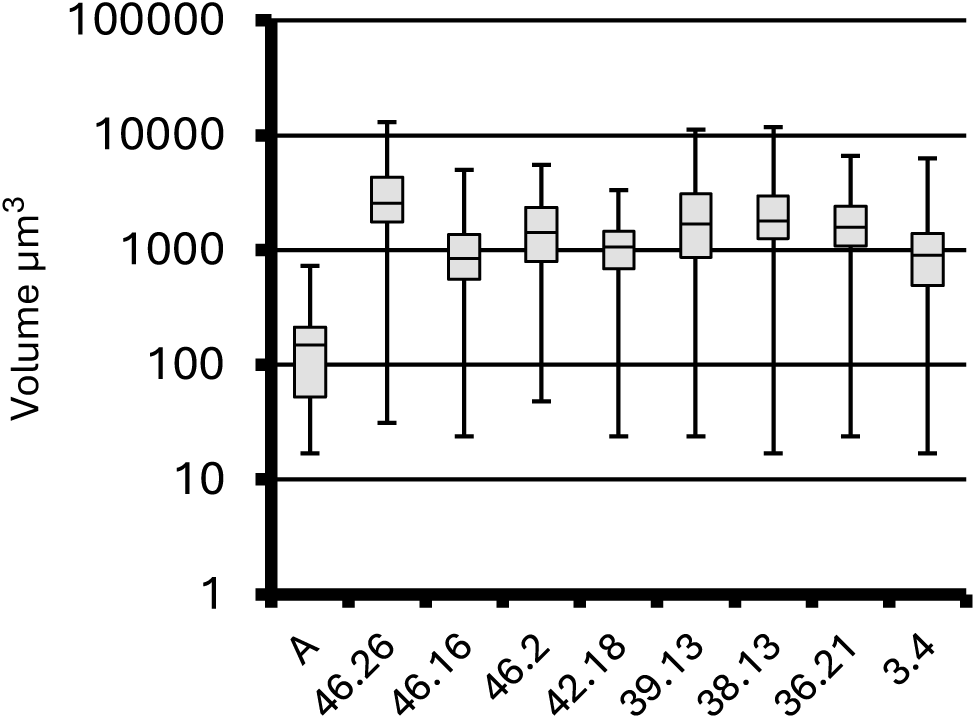
MC isolate cultures’ S50 (the center of biomass) is consistently an order of magnitude greater than the UC ancestor. Boxplots of multiple independent MC isolates (46.26. 46.16, 46.2, 42.18, 39.13, 38.13, 36.21, 3.4) compared to UC ancestor (A, DB146). The middle bar in each grey rectangle represents that culture’s S50. The lower and upper box bounds are S75 and S25 (points in a volume distribution where 25% or 75%, respectively, of biomass is contained in clusters larger than these volumes). Whiskers span the entire population. S50 values of all MC cultures were significantly different from S50 of the unicellular ancestor (Mann-Whitney test, *p*-value <0.001 in all cases).

### Meiotic purification followed by genomic analysis reveals the genetic basis for multicellularity in wild yeast strain DB146

Because the MC phenotype segregated as a Mendelian trait, we set out to find the locus responsible for the MC phenotype in haploid derivatives of DB146, first by using complementation and then by candidate gene sequencing. Because these approaches proved unsuccessful, we turned to meiotic purification (e.g., (39)) to discover the underlying basis for multicellularity. This approach consisted of backcrossing DB146 haploids displaying the multicellular phenotype with a haploid unicellular reference strain, BY4742, then repeatedly backcrossing haploid multicellular progeny from such matings into the reference strain (**Figure 4**). We reasoned that this approach would enable us to pinpoint the multicellularity locus as a genomic region introgressed from the initial multicellular strain.

**Figure 4.**
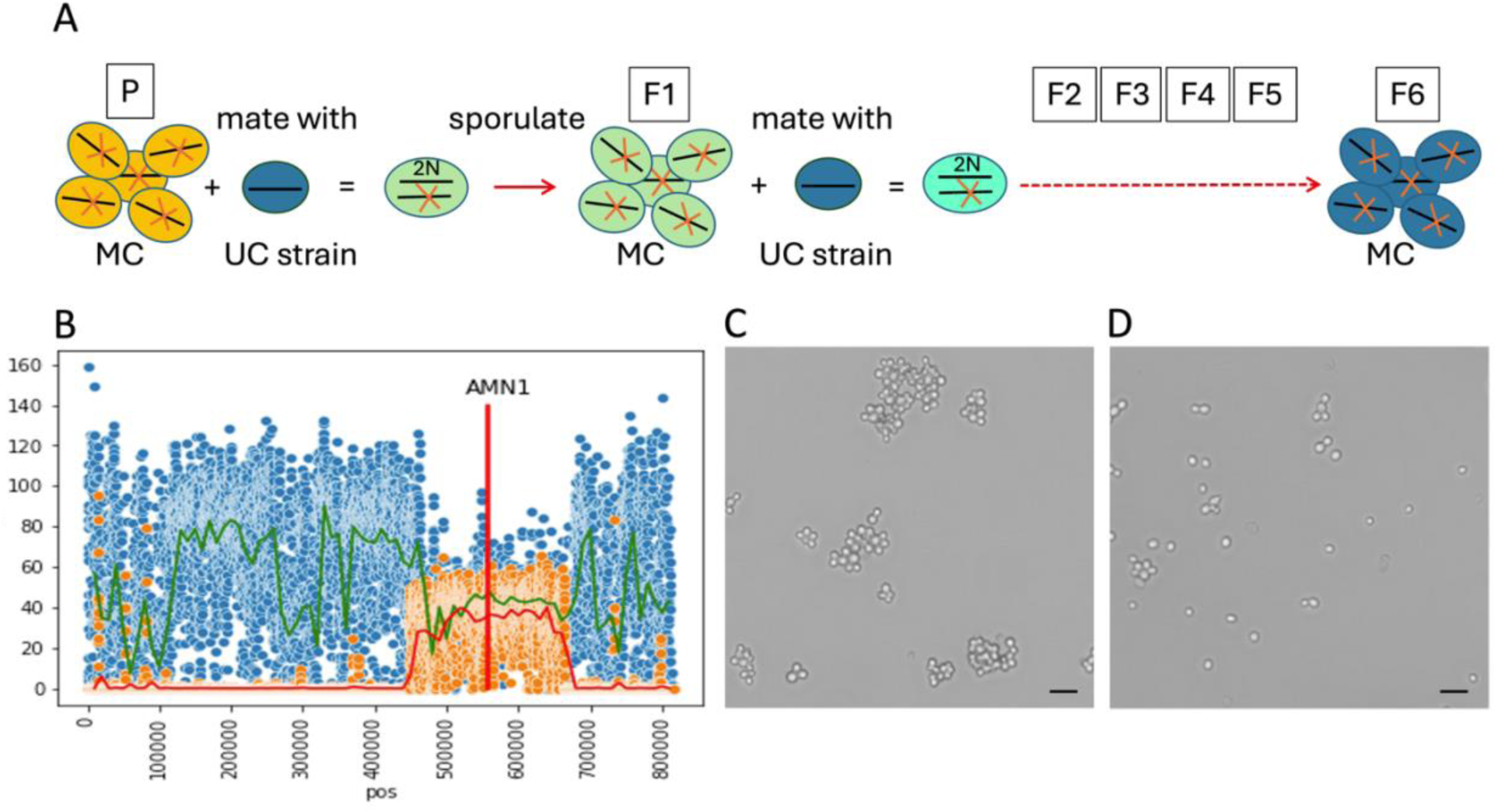
Finding the genetic determinant of multicellularity in haploid derivatives of DB146. **A**: Schematic of meiotic purification to isolate the genomic region carrying the determinant for the multicellular phenotype. P is an MC derivative; F1-F6 are products of successive backcrosses with unicellular lab strain BY4742. **B**: The introgressed region on Chromosome II in the F6 progeny carries the locus for *AMN1.* **C:** Photomicrograph of an MC isolate *before* and **D:** *after* the native *AMN1*DB146 was replaced with the orthologous sequence from a laboratory strain BY4742. Scale bar length = 20μm

In all, we performed 6 rounds of meiotic purification using MC7, a multicellular haploid derivative of DB146, and the unicellular haploid of opposite mating type BY4742 ((40) and (**Figure 4**)). Diploid hybrids produced from the cross were unicellular, as expected. These hybrids were then sporulated and subjected to random spore analysis. Several MC haploid isolates were identified, tested for mating type using mating type PCR, and then mated with BY4742 again. Multicellular offspring were mated with BY4742 in this manner for four additional crosses to yield the MC F6 progeny (**Figure 4A**).

Four independent F6 MC clones were sequenced, and the DB146 introgressed regions were determined by read coverage against the S288c and DB146 genomes (**Figure 4B**). A ∼200 kb region from the DB146 ancestor was found to be present in all four isolates on chromosome II. This introgressed region contained *AMN1*, a gene governing post-mitotic cell separation that has been previously implicated in yeast multicellularity (19). Moreover, the sequence of the *AMN1*^DB146^ allele was identical to the *AMN1*^368D^ variant previously described as the cause of the clustering phenotype in the natural yeast strain YL1C (19). Similar to DB146, only haploid progeny of YL1C display the clustering phenotype (18). Only one-half of DB146 haploid progeny was multicellular because DB146 is hemizygous for the wild-type HO allele (22). Following sporulation, DB146 haploid progeny having a functional HO allele auto-diploidize and retain the unicellular phenotype.

To prove that *AMN1*^DB146/368D^ caused the multicellular phenotype, we replaced it with an allele obtained from laboratory strain BY4742. Several MC haploid DB146 spore isolates were transformed with a linear construct containing the *amn1*^BY4742^ marked with the g418 resistance marker flanked by regions homologous to the regions upstream and downstream of *AMN1* in DB146. The allele replacement *AMN1*^DB146^ → *amn1*^BY4742^ resulted in a unicellular phenotype (**Figure 4: C,D**), supporting our claim that *AMN1*^DB146^ is indeed the locus causing multicellularity in DB146’s haploid derivatives.

### Multicellular cell clusters grow more slowly but live longer than single cells

To understand why the MC phenotype was retained in wild strain DB146, we evaluated its reproductive capacity and survivorship relative to that of its UC ancestor. In the wild, a genotype’s success in contributing progeny to future generations depends on its rate of increase and survival, especially in relation to other genotypes of the same species living in sympatry. To assess reproductive capacity, we estimated two standard growth parameters for 8 MC strains and their UC ancestor DB146, grown at 24°C in rich medium containing 1% glucose: maximum specific growth rate (µ_max_) and yield at stationary phase (Y_s_). UC ancestor DB146 was found to grow faster and to a higher yield than all MC strains that were its meiotic progeny (UC µ_max_=0.59±0.01 hr^-1^ vs. MC µ_max_=0.49±0.02 hr^-1^, UC Y_s_=1.51±0.01 vs. MC Y_s_=1.26±0.03, in OD600 units). Thus, clustered MC derivatives should not have an advantage over their UC ancestor when the yeasts grow in liquid batch culture under optimal conditions.

Next, we tested the survivorship of clusters and unicells under strongly adverse conditions: 100 days of complete starvation. Given that each MC cluster (or ‘body’) is clonal, and that any viable cell in a cluster can convey that clone’s genotype into the next generation, we decided to score any cluster with at least one viable cell remaining as “viable.” The viability of 5 MC isolates and their DB146 ancestor was analyzed in triplicate. Cells were stained, photographed and counted every 10 days for 100 days. At each time point, at least 200 individual clusters or cells were stained with Trypan blue exclusion dye and counted. Individual cell viability was similar for single cells and cells in clusters; however, as starvation progressed beyond 40 days, the viability of MC clusters (bodies) surpassed that of the UC control (**Figure 5**). By 100 days, the average viability of individual MC clusters was >85%, whereas that of the UC ancestor was ca. 60%, a statistically significant difference (p-value < 0.05, Student t-test).

**Figure 5.**
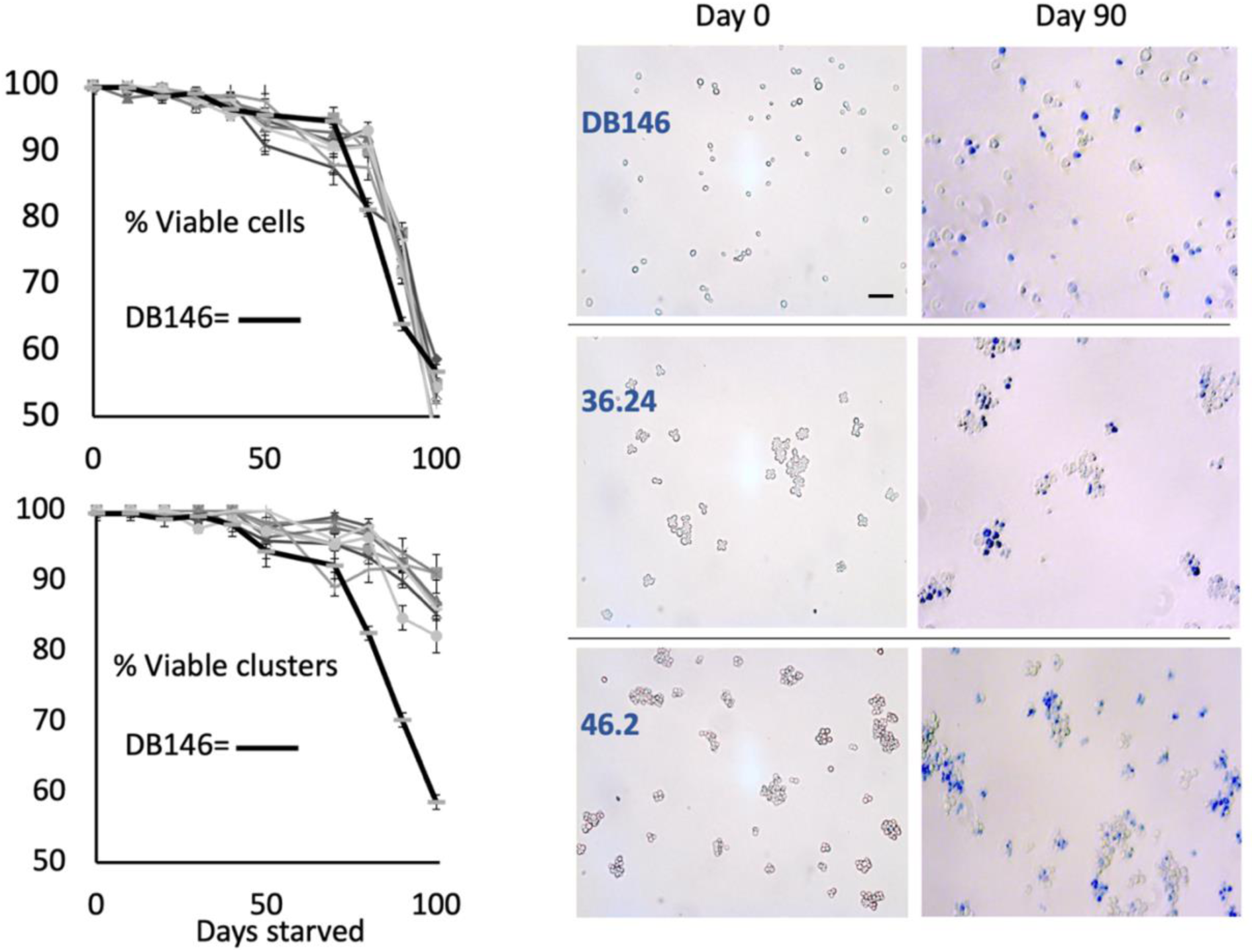
Cell and cluster viability during 100-day starvation experiments. ***Upper graph***: the dynamics of survivorship of *individual cells* is indistinguishable between diploid DB146 and its haploid MC derivatives. ***Lower graph:*** By contrast, when MC cells are considered components of a ‘body’ (see text), bodies consisting of MC derivatives survive better than bodies of their UC DB146 ancestor. ***Photomicrographs***: Trypan blue staining of the UC (DB146) culture and two representative MC cultures (36.24 and 46.2) at day 0 (100% viability) and after 90 days of starvation. Scale bar length = 20μm

In summary, all MC isolates grew more slowly and to a lower overall cell density than did UC isolates. Over time, however, survivorship became significantly higher per body basis in MC strains than in the UC ancestor, providing a new, group-level Darwinian entity (41). Cryptic multicellularity concealed in the hemithallic DB146 genome lacks obvious adaptive value when cells proliferate under favorable conditions. However, yeast cells in nature are subject to “boom and bust” cycles in nutrient availability, and “bust” is often a signal to initiate sporulation. Our starvation data suggest that uncovering cryptic multicellularity in DB146’s haploid meiotic progeny could prove adaptive under slow-growth or non-growth conditions. To explore this possibility, we analyzed the fitness of MC strains versus UC strains in pairwise (head-to-head) competitions by subjecting cells to stressful conditions likely to enforce slow growth or no growth at all.

### Pairwise competition assays show that the cryptic multicellular phenotype can be selectively advantageous under adverse conditions

If the capacity to form clusters is preserved in the genomes of wild yeast, then that capacity should be beneficial under at least some set of environmental conditions encountered by wild yeast in nature. To ascertain what these conditions might be, we competed multicellular, haploid yeast (MC) against their unicellular, diploid ancestor (UC), as these phenotypes would compete against each other in the wild. Specifically, DB146 diploids that underwent meiosis and sporulated would produce two phenotypes: unicells originating from spores that diploidized and clusters originating from spores that did not. Our experiments focused on slow- or non-growing cells under conditions that differed in nutrient and water availability, temperature, initial cell density, and combinations thereof, i.e., conditions that cells would likely encounter in the wild.

To enumerate the relative abundance of MC and UC phenotypes in co-culture, we used direct plating rather than a particle analyzer (for details, see **Methods**), reasoning that particle volume might inaccurately represent total cell count per cluster due to irregularities of cluster packing. MC colonies can be easily distinguished from UC colonies based on colony morphology. Instead of the smooth colony surface produced by UC cells, MC clusters produce colonies that exhibit ruffled edges, dimpled centers, and/or ridges (see **Supplementary Materials: Figure S1)**. We were particularly interested in the survival of a cluster versus a single cell, because if even one cell in a cluster is alive at the end of a competition experiment, the ‘body’ formed by that cluster should still be considered ‘alive’.

Competition under strongly adverse conditions is often a game of survival, where the viabilities of competing individuals decrease and populations contract, rather than expand. To evaluate the outcome of such competitions, we used the selection rate constant per day (42). In a survival competition, the Malthusian parameters of competing populations become negative. However, this does not negate the relevance of the resulting selection rate. To calculate selection rates in our competitions, we made the assumptions of weak selection, no mutation, and a constant carrying capacity close to zero. To arrive at an average selection-rate constant for the MC phenotype, we ascertained whether the data from different MC derivatives (specifically, 44.26, 46.2, 46.16, and MC7) were similar before averaging them across the MC phenotype. We compared colony counts for all time points (15, 30, and 45 days post-inoculation), normalized to Day 0 colony count for all MC derivatives in each competition. We confirmed that these data were not significantly different (*p*-values for all experiments >>0.05, one-way ANOVA). This allowed us to average the selection rates for all MC derivatives.

We first set up competitions at a high titer of 5 × 10^7^ cells mL^-1^. Mixed populations of unicells and clusters were incubated at 24°C in 1% YPD (see Methods). As cells tend to settle in the tube, effectively changing their local density, test tubes were slowly shaken at 76 rpm to keep cells in suspension. In these experiments, MC showed no significant advantage over UC over a 45-day time course (mean selection rate over three time points of 15, 30 and 45 days 4.77 × 10^-3^ ± 3.33 × 10^-3^, **Figure 6**: the first bar – high titer, high temp, high nutrients). Interestingly, under these conditions, the calculated selection rate over the first 15 days was negative for MC cultures, whereas over the second (15-30 days) and the third (30-45 days), selection rates were slightly positive for MC cultures, reflecting the onset of starvation. Indeed, under prolonged starvation, MC cultures exhibit an advantage, as was shown in the previous section.

**Figure 6.**
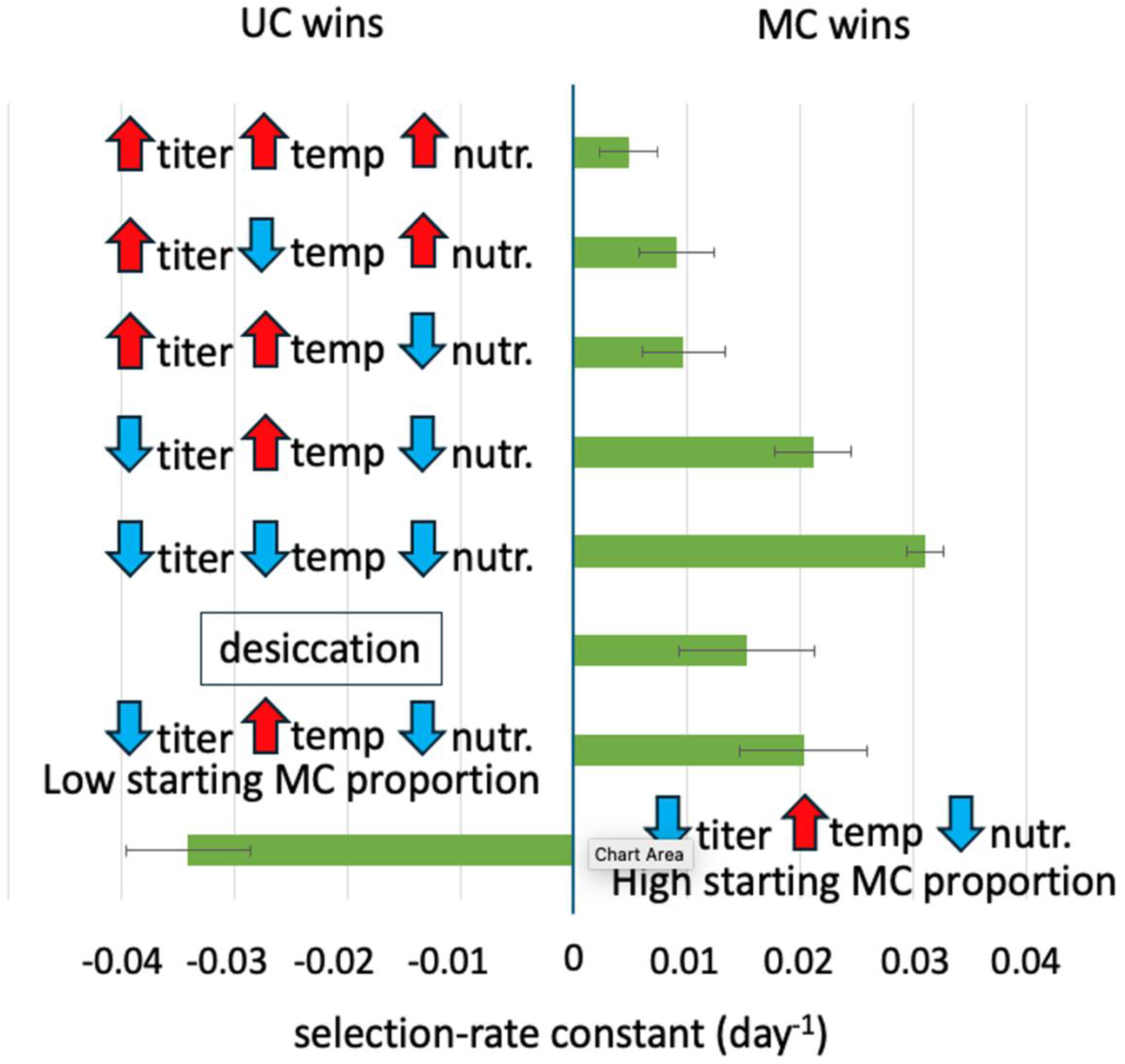
MC isolates have a survival advantage under adverse conditions. Bars represent mean selection-rate constants (1/day) for cultures in which MC were competed head-to-head against UC. Error bars are SEM of selection rates obtained between 0 to 15, 15 to 30 and 30 to 45-day periods. **Red** and **blue** arrows denote, respectively, denote **high** and **low** values for initial cell density, temperature, or nutrient level (see text for details). Except where noted, MC and UC cultures were mixed in 1:1 proportion. Low and high starting proportions were 1:10 or 10:1 MC:UC, respectively.

Next, we decided to explore conditions reminiscent of those yeast populations might experience in the wild. Low temperatures, nutrient scarcity, low local cell density and even periodic desiccation are all part and parcel of the existence of temperate zone yeast in their natural environment.

Accordingly, we carried out MC *versus* UC competitions at low temperature (10°C), where MC cultures showed an advantage (selection-rate constant of 9.11 × 10^-3^ ± 3.70 × 10^-3^ day^-1^) (**Figure 6**, the second bar), and in distilled water (low nutrients), where MC cultures also showed an advantage (selection-rate constant of 9.68 × 10^-3^ ± 3.37 × 10^-3^ day^-1^) (**Figure 6**, the third bar). Next, we carried out MC *vs.* UC competitions at low titer (10^4^ cells mL^-1^) using distilled water instead of rich medium, reasoning that in the wild, yeast cells might persist at low density under very low-nutrient conditions. Cultures were incubated at either 24°C or at 10°C, the latter to simulate conditions cells might experience overwintering. Under these conditions, MC haploid progeny showed an even higher selective advantage over their UC ancestor (mean selection-rate constant of 2.32 × 10^-2^ ± 0.07 × 10^-2^ day^-1^ at 24°C, and 3.11 × 10^-2^ ± 0.39 × 10^-2^ day^-1^ at 10°C) (**Figure 6**, the fourth and fifth bars, respectively).

Because another variable likely to be encountered by yeast in nature is a change in water availability, we also subjected a subset of competing cultures to periodic desiccation at high temperature, high titer and high nutrients. Desiccation alone uncovered a significant selective advantage for MC strains over their UC ancestor (selection-rate constant of 1.52 × 10^-2^ ± 0.56 × 10^-2^ day^-1^) (**Figure 6**, the sixth bar). However, it should be noted that desiccation significantly decreased overall viability for both phenotypes as judged by the “pure culture” controls (data not shown). Finally, we tested for the effect of different ecological “starting points” by competing the UC ancestor and its MC derivatives at different initial proportions in ddH_2_O at 24°C. When MC was initially in the minority (1:10 MC:UC ratio), the MC phenotype prevailed (selection-rate constant of 2.03 × 10^-2^± 0.39 × 10^-2^ day^-1^) (**Figure 6**, the seventh bar), whereas when competitions were initiated at a 10:1 MC:UC ratio, the MC phenotype was at a competitive disadvantage (selection-rate constant of -3.42 × 10^-2^ ± 0.89 × 10^-2^ day^-1^) (**Figure 6**, the eighth bar).

To summarize our results from two sections related to growth and competition, our data show that the conditions under which the unicellular phenotype prevails or is not outcompeted, are: elevated temperature, high initial nutrient levels, even with subsequent nutrient depletion during the period of competition, high cell density, or low initial proportion of UC cells. By contrast, conditions under which the multicellular phenotype prevails are low temperature, scarce nutrients, low cell density and combinations thereof, as well as cycles of desiccation-rehydration, and a low initial proportion of MC cells. Several conditions under which the MC phenotype proves selectively advantageous are reminiscent of winter in the temperate zone, conditions that wild yeast, e.g., champagne yeast DB146 is likely to encounter in the wild.

### Cryptic multicellularity phenotype is widespread in other wild Saccharomyces strains

If the multicellular phenotype is adaptive under different adverse conditions, then we would expect to find that phenotype in wild yeast strains besides DB146. To test this expectation, we screened 19 wild isolates of *S. cerevisiae* and one lab strain (BY4743) for multicellularity (**Table S1**). The sources from which these 19 strains were isolated included vineyards, fruit, and oak tree bark and exudate. The original diploid strains and their meiotic progeny, derived by random spore analysis or tetrad dissection, were genotyped for ploidy by mating-type PCR (see Materials and Methods) and examined by light microscopy. Strains and derivatives that appeared to be multicellular were treated as described with EDTA/sonication and with zymolyase to distinguish between strains that exhibited aggregative multicellularity by flocculation and those that exhibited clonal multicellularity because daughter cells did not separate from their mothers. Only those isolates that exhibited clonal multicellularity were scored as multicellular.

Of 19 wild strains, which were all diploid, 6 strains (EC-1118, 8130, E85, E684, PM229 and PM264) were found to be multicellular or to give rise to multicellular progeny (Fig. 7). Of these strains, two came from vineyards, two from fruit, and two from unknown sources. PM264 displayed the MC phenotype, and its meiotic derivatives were also MC. In contrast, PM229 was UC, all meiotic derivatives – diploid, and approximately half of the meiotic derivatives were multicellular. For strains EC-1118, 8130, E85 and E684, their diploid progeny was UC and haploid progeny – MC. To determine whether multicellularity in the haploid offspring of EC-1118, 8130, E85 and E684 was due to their ploidy, the haploid meiotic progeny of these strains were transformed with a plasmid carrying a copy of the HO endonuclease gene under the control of its endogenous promoter (pHS2) (Addgene # 81037). Upon autodiploidization, all MC offspring of these strains reverted to the UC phenotype. This suggests that the MC phenotype in haploid derivatives of these 4 strains may be due to a mechanism similar to that we uncovered in DB146, whereas the nature of multicellularity for PM264 and PM229 is presently unknown.

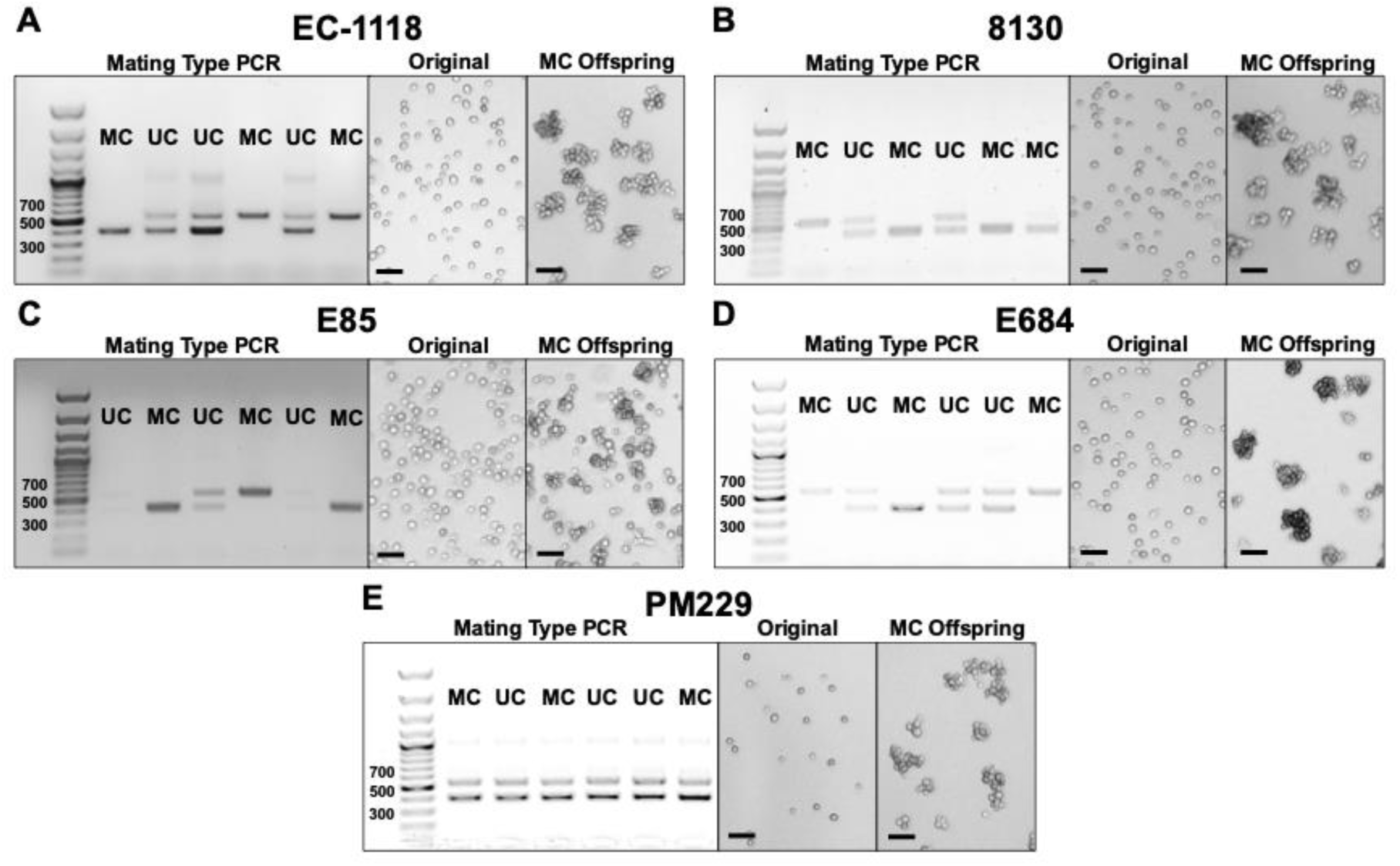
**Supplemental Figure X.** Ploidy and clustering phenotypes of wild yeast offspring. Multicellular (MC) progeny were derived from unicellular (UC) wild yeast strains (A) EC-1118, (B) 8130, (C) E85, (D) E684, and (E) PM229. For each strain, the results from mating type PCR on progeny from tetrad dissection are displayed along with micrographs of the original strain and a representative MC offspring. For mating type PCR, the presence of both a 404 bp band and a 544 bp band indicates that the offspring is diploid while the presence of only one of these bands indicates that it is haploid. Scale bar length = 20 μm.

Overall, a substantial proportion (20%) of this modest sample of wild *Saccharomyces* strains exhibits multicellular phenotypes that can be attributed to genetic mechanisms that are either ploidy-dependent or ploidy-independent, further supporting the view that clonal multicellularity can be adaptive in wild yeast. This estimate may be conservative, as a homothallic yeast strain that carries the ploidy-driven multicellularity trait via *AMN1*^368D^ will only transiently show multicellularity and then convert to UC after mating–type switching and diploidization.

## Discussion

The advent of multicellularity is widely viewed as a major evolutionary transition in the history of life, opening the door not only to large size but also to the possibility of division of labor via cellular differentiation. Multicellularity has independently arisen dozens of times, beginning 3 billion years ago with the cyanobacteria (43). However, in most instances, *de novo* multicellularity did not lead to ‘complex multicellularity’ in the sense of large, relatively long-lived organisms composed of many cell types (44). This is especially true of the many cases where simple multicellularity has arisen in prokaryotes (45). In recent years, considerable attention has been focused on eukaryotic lineages that gave rise to “complex multicellularity,” especially those having extant representatives, some of which are unicellular, others of which are multicellular. Of particular interest in this regard are protists like choanoflagellates, which are phylogenetically affiliated with animals (46), *Chlamydomonas* spp., which are classified among the green algae (47), and *Saccharomyces* spp., which are ascomycete fungi (48). Simple, heritable multicellularity has been repeatedly evolved in the laboratory from unicellular *Chlamydomonas reinhardtii* (16,49) and *Saccharomyces cerevisiae* (2,14). In both species, the experimental end result is clonal multicellularity, where each cluster originates from a single genotype rather than from an aggregate of possibly different genotypes. Also, in both species, the experimental end result is obligate multicellularity, in which the evolved phenotypes are committed to a predominantly multicellular life history, with unicells or cluster fragments serving as propagules.

### Cryptic multicellularity in champagne yeast DB146 is ploidy-dependent

Here, we investigated facultative multicellularity in wild *Saccharomyces cerevisiae*, focusing on genetic and environmental influences that favor its expression. In our investigation, multicellularity proved to be a Mendelian trait whose expression depends on unveiling a mutant version of the *AMN1* gene in haploids. *AMN1* encodes a Ste12p-regulated E3 ubiquitin ligase that modulates post-mitotic mother-daughter cell separation and helps to reset the cell cycle in G1 (19,50). The *AMN1* mutant we recovered in wild yeast strain DB146 is defective for cell separation in haploids, resulting in clonal clusters in which cells are tethered to one another by their cell walls. This finding is reminiscent of how clonal multicellularity can arise in evolving lab yeast strains via mutation of the transcriptional regulator, *ACE2,* which is also involved in cell cycle control and control of mother-daughter cell separation (5,51). Indeed, Amn1p post-translationally controls Ace2p degradation via the yeast ubiquitin proteasome system, thereby connecting the RAM (regulation of <u>A</u>ce2 and <u>m</u>orphogenesis) network with that of the MEN (<u>m</u>itotic <u>e</u>xit <u>n</u>etwork) (19).

Our finding of ploidy-dependent facultative multicellularity in the champagne strain DB146 is consistent with the recent findings of Barrere et al. (21), who used flow cytometry to score the clustering phenotypes of 22 phylogenetically diverse wild yeast from the NCYC SGRP2 collection (52). Haploid isolates of both mating types exhibited significantly greater clustering scores than their diploid progenitors, indicating that in wild yeast, facultative multicellularity is at least partly under the control of the mating type locus. Ploidy-dependent facultative multicellularity in wild yeast isolates is reminiscent of the evolution of bistable life cycles with multilevel fitness advantages in *Pseudomonas* (53), though the wild yeast described here assumes a multicellular or unicellular state with the life-cycle switch dependent on ploidy rather than on mutation, as in (53).

### Advantages and disadvantages of cryptic, AMN1-driven multicellularity: Stress resistance

Specific *AMN1* mutations underpin the formation of multicellular clusters in *Saccharomyces cerevisiae*, while others reverse this phenotype, leading heritably multicellular yeast to become unicellular. Kuzdal-Fick et al. (7) used this as a tool to explore the costs and benefits of experimentally evolved unicellularity. Ancestral multicellular forms were found to be more resistant to freeze-thaw, H_2_O_2_, and ethanol stress than their unicellular descendants; however, stress-resistance traded off against growth rate and yield upon entry to stationary phase, which were higher in unicells. Similarly, we observed that under conditions such as low temperature and starvation, multicellular DB146 haploids generally showed higher survivorship than their unicellular ancestor, provided that both unicells and clonal clusters are regarded as ‘bodies’. These outcomes are reminiscent of the latent potential for multicellularity among certain unicellular green algae. For example, transient multicellularity can be induced in unicellular *Chlamydomonas* spp. by environmental stressors such as low ambient Ca^2+^ or high PO_4_ concentrations (49), either of which leads to palmelloids wherein multiple daughter cells are retained within a shared extracellular matrix. These multicellular forms demonstrate a fitness advantage over their unicellular progenitors under stressful conditions like high light intensity (54,55) or predation (16,56). However, under conditions where resources are non-limiting, unicellular *Chlamydomonas* grow more rapidly and to higher cell densities (8).

### Advantages and disadvantages of cryptic multicellularity: Famine and feast

When temperate zone yeast overwinters, dead and dying cells may become the only source of nutrients needed to sustain cells that are still alive. Indeed, recent work has shown that cell cannibalism in multicellular bodies (also called cell-in-cell) is found across the Tree of Life, and that specific cell-in-cell genes are associated with normal development as well as with cancer (57). In dead or dying cells, such nutrients are chiefly stored as complex polymers that require extracellular digestion prior to assimilation. Like all fungi, *S. cerevisiae* can access complex carbohydrates, proteins, and lipid-like molecules by exporting a variety of hydrolytic enzymes (58–62). These secreted glucanases, proteases, and lipases become “public goods” whose effective concentration, and therefore activity, is greater in dense multicellular clusters than in single cells. Indeed, Murray and colleagues have shown that scarce nutrients that require extracellular processing are more readily extracted from the environment by cell clusters than by single cells, especially at low titers of each cell type (18). In our experiments, these physiological advantages are realized as increased Darwinian fitness when multicellular haploid yeast compete head-to-head with unicellular haploids. Significantly, these experiments were carried out under prolonged starvation. Thus, we speculate that in haploid *AMN1* mutant yeast, multicellularity may carry a two-fold advantage: cells in clusters build up more public goods enzymes, and cells in clusters act as a resource for one another. Whether or not there is a pattern or program underlying which cells in a cluster die, as well as when they die, remains to be discovered.

Our data show that while the cluster-forming trait is disadvantageous under optimal growth conditions, it can be beneficial under slow growth or non-growth conditions. Multiple sub-optimal conditions favored the survival of multicellular clusters over unicellular strains: low temperature, nutrient scarcity, low density, desiccation and low initial proportion of multicellular ‘bodies.’ Taken together, these typify winter conditions faced by wild yeast living in the temperate zone. During winter and periods of famine, it is survival that defines fitness – until the spring and summer feast, when haploid cells in a cluster may mate and resume a diploid unicellular lifestyle that allows for prolific growth under conditions where temperature and humidity are favorable, and nutrients are abundant.

### Cryptic multicellularity is not uncommon in wild yeasts

To ascertain whether the multicellular phenotype is found among other wild *Saccharomyces*, we examined yeast isolated from diverse habitats. We found that at least 20% of these strains exhibit clonal multicellularity and that the phenotype can be driven by different genetic mechanisms, which include ploidy-dependent unmasking of *AMN1*. This genetic flexibility adds to the diversity of mechanisms that enable yeast to become multicellular. Some of these result in aggregative rather than clonal multicellularity owing to a mutation in *FLO1* and its regulators (4). Others result in pseudohyphal growth arising from regulatory changes in the expression of *MUC1/FLO11* triggered by activation of the MAP kinase cascade (63). Significantly, both these types of yeast multicellularity are triggered by nutrient limitation, particularly in the forms of nitrogen and/or carbon. Overall, we find that while the unicellular phenotype shows clear advantages under optimal growth conditions in the lab, cryptic multicellularity can be selectively advantageous under adverse conditions that cells frequently encounter in the wild. In this view, alleles of *AMN1* that cause a defect in cell separation may be an evolutionary adaptation that allows haploid cells to survive during overwintering. From this perspective, *AMN1* alleles derived from strictly unicellular laboratory strains may have been inadvertently selected for the ease of cell manipulation in the laboratory, and the *AMN1*^368D^ allele that causes facultative multicellularity is indeed the *bona fide* wild-type. Nascent multicellularity, especially in a heterogonic species that undergoes mating-type switching, could be maintained under balancing selection by cycles of famine and feast, cold and warm, dry and wet, until circumstances arose that favor the complete unmasking of this trait and the evolution of cellular differentiation.

## Acknowledgments

The authors are grateful to Emily Cook for technical assistance, to William Ratcliff, Peter L. Conlin, Matthew Herron and Francesca Storici for insightful comments and suggestions, and to our reviewers, who significantly improved the manuscript.

## Data Availability Statement

The authors confirm that all data necessary to support the conclusions of the article are presented within the article, figures, and tables. Strains and plasmids are available upon request.

## Supplementary Materials: Methods

### Cloning of the MC allele

Since DB146 only expresses a multicellular phenotype in the haploid state, we hypothesized that a haploinsufficient determinant was responsible for the haploid MC phenotype. We tested this by transforming the multicellular haploid derivative MC7 with a *S. cerevisiae* 2µ-based genomic library (65). To enrich for the UC phenotype, the population of 4×10^4^ of library transformants was allowed to partially settle in a test tube. Transformants were sampled from the top of the suspension, checked for unicellularity and then serially passed on rich medium to cure genomic library plasmids and observe the reversion to the original multicellular phenotype. No reversions to the multicellular phenotype were observed, suggesting that the library plasmids were not responsible for the reduction in multicellularity.

### Candidate gene sequencing

Since the MC phenotype segregates as a Mendelian trait, we sought to find it by sequencing. To determine genetic candidates, we sequenced both a diploid unicellular form of DB146 and several multicellular spore derivatives of DB146. Nonsynonymous SNPs were determined and then compared to several genes known to cause a multicellular phenotype in yeast, namely *ACE2*, *CTS1*, *AMN1* and *SWI5* (50,66–68) (**Table S2**). Of these four loci, only *ACE2* lacked any nonsynonymous mutations.

#### Determination of MC or UC origin of colonies by their appearance

##### Cluster analysis on a particle analyzer

Here, we used a particle counter (Multisizer 4e, Beckman Coulter Inc., Brea, CA) to resolve different size distributions. Overnight cultures of DB146 derivatives (multicellular and unicellular) were examined on a hemocytometer to determine their titer. MC and UC suspensions were then mixed to produce cell suspensions that had different percentages of clusters versus unicells, (0%, 5%, 10%, 15%, 20%, 40%, 45%, 50%, 55%, and 60%); these were run in triplicate on the Coulter Counter (**Supplementary Materials: Figure S2**). The resulting data were converted to their respective S50 values and graphed as related to their respective MC-UC mixtures. This experiment shows that particle counting to distinguish different proportions of clusters vs. single cells is not particularly sensitive to a change in cluster/unicell ratio, especially in 40-60% mixtures.

---

**Figure S1.**
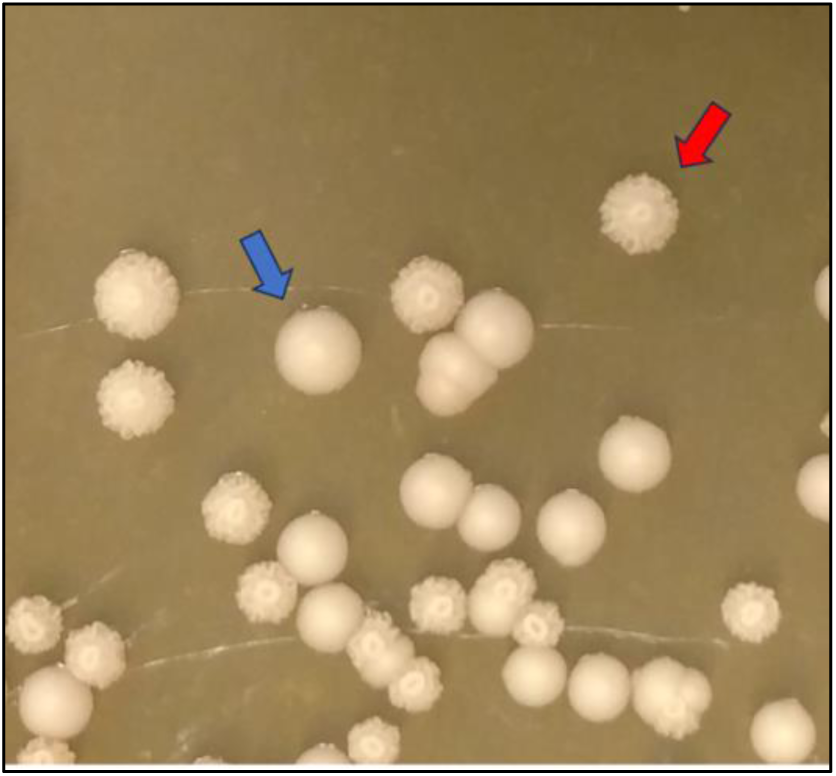
Plating of a mixture of MC and UC cultures. Single cells form smooth colonies (blue arrow). MC clusters form colonies with a wrinkled surface (red arrow), allowing simple identification.

**Figure S2.**
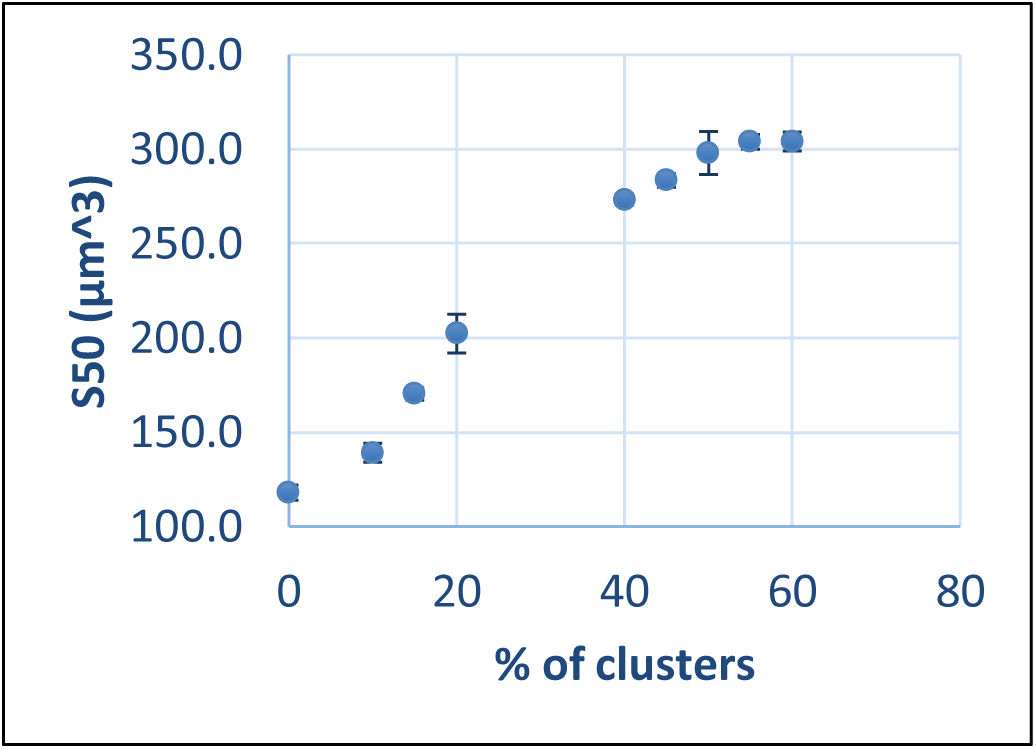
Sensitivity of S50 to the proportion of MC in mixed multicell and unicell populations. X-axis is the proportion of MC in the MC-UC mixed population; Y-axis is S50 in cluster volume, μm3, bars are SEM.

**Table S1.**
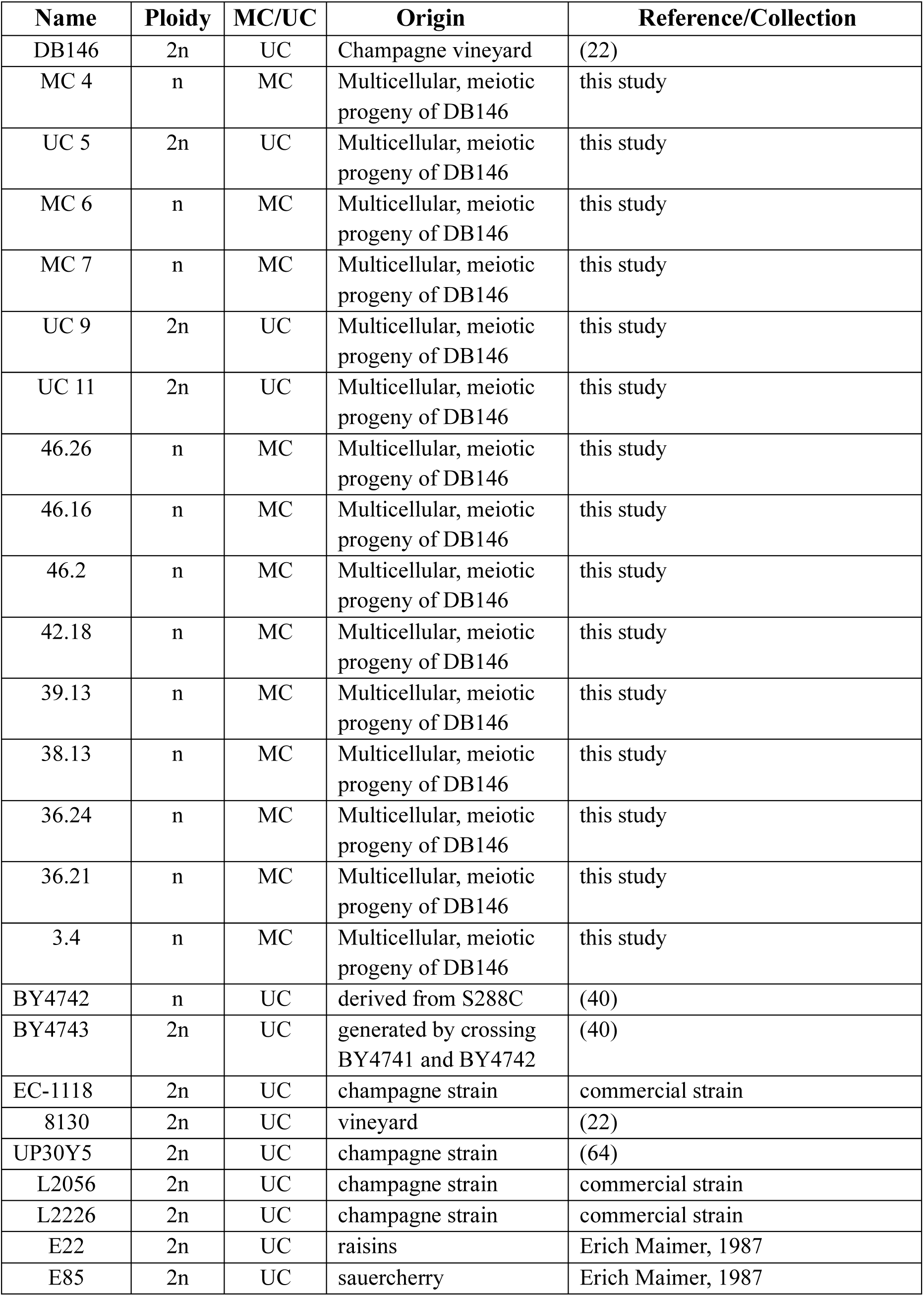
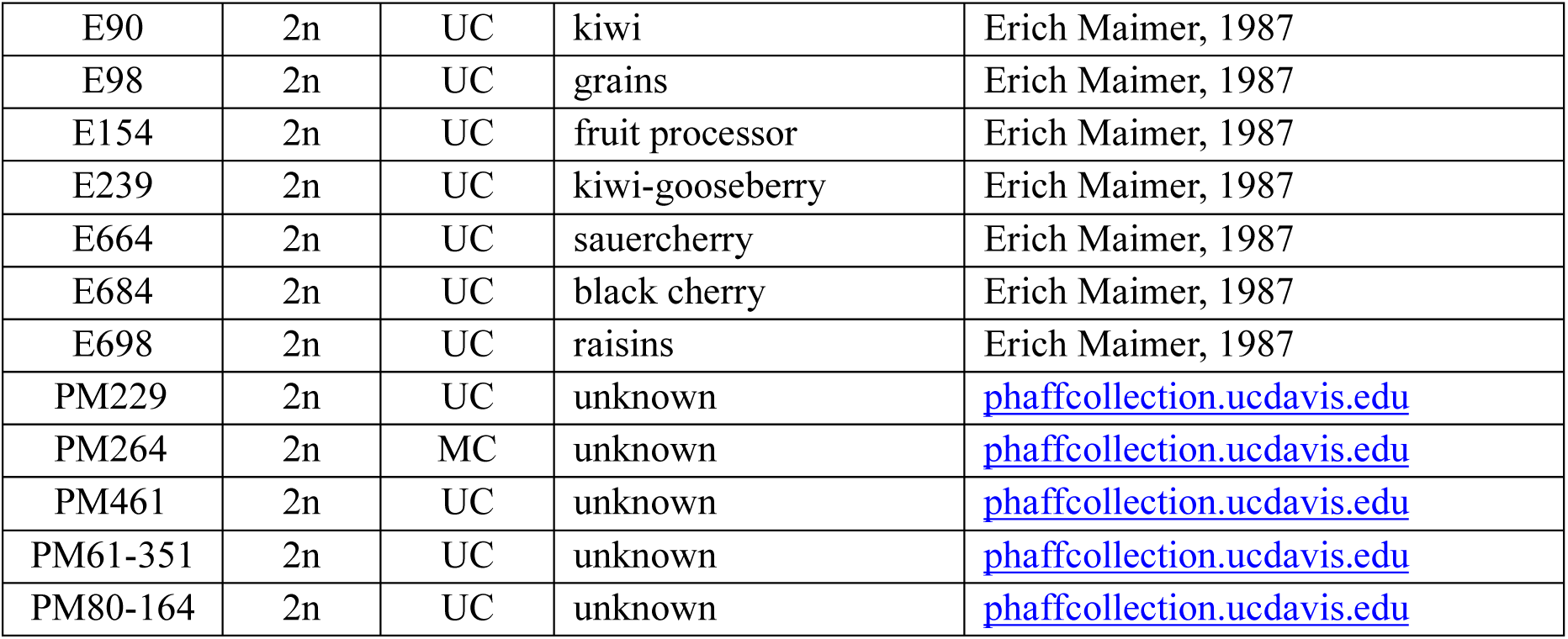
Strains used in this study.

**Table S2.** Nonsynonymous mutations in genes reported to cause an MC phenotype. Genes known to cause a multicellular phenotype were screened by alignment of DB146 reads to the S288c genome and translation of the reading frame. Nonsynonymous mutations were observed at the *AMN1*, *CTS1* and *SWI5* loci, providing several candidate loci that might explain the multicellular phenotype. Therefore, we did not find the single locus that causes multicellularity in DB146 haploid derivatives using the candidate gene sequencing approach.

|  |  |
| --- | --- |
| AMN1 | 226 R -> Q |
| AMN1 | 233 H->R |
| CTS1 | 16 L->P |
| CTS1 | 23 R->S |
| CTS1 | 321 T->S |
| CTS1 | 327-331 Deletion |
| CTS1 | 433 P->S |
| CTS1 | 434 I->L |
| CTS1 | 468-472 Deletion |
| SWI5 | 16 L-> Q |
| SWI5 | 494 V->I |

## References

1. Pentz JT, MacGillivray K, DuBose JG, Conlin PL, Reinhardt E, Libby E, et al. Evolutionary consequences of nascent multicellular life cycles. Elife. 2023 Oct 27;12:e84336.

2. Bozdag GO, Zamani-Dahaj SA, Day TC, Kahn PC, Burnetti AJ, Lac DT, et al. De novo evolution of macroscopic multicellularity. Nature. 2023 May;617(7962):747–54.

3. Márquez-Zacarías P, Conlin PL, Tong K, Pentz JT, Ratcliff WC. Why have aggregative multicellular organisms stayed simple? Curr Genet. 2021 Dec;67(6):871–6.

4. Hope EA, Amorosi CJ, Miller AW, Dang K, Heil CS, Dunham MJ. Experimental Evolution Reveals Favored Adaptive Routes to Cell Aggregation in Yeast. Genetics. 2017 June;206(2):1153–67.

5. Ratcliff WC, Fankhauser JD, Rogers DW, Greig D, Travisano M. Origins of multicellular evolvability in snowflake yeast. Nature Communications. 2015 Jan 20;6:6102.

6. Pfeiffer T, Bonhoeffer S. An evolutionary scenario for the transition to undifferentiated multicellularity. Proceedings of the National Academy of Sciences of the United States of America. 2003 Feb 4;100(3):1095–8.

7. Kuzdzal-Fick JJ, Chen L, Balázsi G. Disadvantages and benefits of evolved unicellularity versus multicellularity in budding yeast. Ecol Evol. 2019 July 9;9(15):8509–23.

8. Cornwallis CK, Svensson-Coelho M, Lindh M, Li Q, Stábile F, Hansson LA, et al. Single-cell adaptations shape evolutionary transitions to multicellularity in green algae. Nat Ecol Evol. 2023 June;7(6):889–902.

9. Conlin PL, Goldsby HJ, Libby E, Skocelas KG, Ratcliff WC, Ofria C, et al. Division of labor promotes the entrenchment of multicellularity [Internet]. bioRxiv; 2023 [cited 2025 Oct 21]. p. 2023.03.15.532780. Available from: https://www.biorxiv.org/content/10.1101/2023.03.15.532780v1

10. Kelly B, Carrizo GE, Edwards-Hicks J, Sanin DE, Stanczak MA, Priesnitz C, et al. Sulfur sequestration promotes multicellularity during nutrient limitation. Nature. 2021 Mar;591(7850):471–6.

11. Strassmann JE, Queller DC. Evolution of cooperation and control of cheating in a social microbe. Proc Natl Acad Sci U S A. 2011 June 28;108 Suppl 2(Suppl 2):10855–62.

12. Fisher RM, Shik JZ, Boomsma JJ. The evolution of multicellular complexity: the role of relatedness and environmental constraints. Proc Biol Sci. 2020 July 29;287(1931):20192963.

13. Isaksson H, Lind P, Libby E. Adaptive evolutionary trajectories in complexity: Transitions between unicellularity and facultative differentiated multicellularity. Proceedings of the National Academy of Sciences. 2025 Jan 28;122(4):e2411692122.

14. Ratcliff WC, Denison RF, Borrello M, Travisano M. Experimental evolution of multicellularity. Proc Natl Acad Sci U S A. 2012 Jan 31;109(5):1595–600.

15. Fisher RM, Bell T, West SA. Multicellular group formation in response to predators in the alga Chlorella vulgaris. J Evol Biol. 2016 Mar;29(3):551–9.

16. Herron MD, Borin JM, Boswell JC, Walker J, Chen ICK, Knox CA, et al. De novo origins of multicellularity in response to predation. Sci Rep. 2019 Feb 20;9(1):2328.

17. Bonforti A, Solé R. Unicellular-multicellular evolutionary branching driven by resource limitations. J R Soc Interface. 2022 June;19(191):20220018.

18. Koschwanez JH, Foster KR, Murray AW. Improved use of a public good selects for the evolution of undifferentiated multicellularity. eLife. 2013 Apr;2013(2):e00367.

19. Fang O, Id XH, Wang L, Jiang N, Yang J, Li B, et al. Amn1 governs post-mitotic cell separation in Saccharomyces cerevisiae. Barsh GS, editor. PLOS Genetics. 2018 Oct 1;14(10):1–20.

20. Gresham D, Ruderfer DM, Pratt SC, Schacherer J, Dunham MJ, Botstein D, et al. Genome-wide detection of polymorphisms at nucleotide resolution with a single DNA microarray. Science. 2006;311(5769):1932–6.

21. Barrere J, Nanda P, Murray AW. Alternating selection for dispersal and multicellularity favors regulated life cycles. Curr Biol. 2023 May 8;33(9):1809–1817.e3.

22. Bidenne C, Blondin B, Dequin S, Vezinhet F. Analysis of the chromosomal DNA polymorphism of wine strains of Saccharomyces cerevisiae. Curr Genet. 1992;22(1):1–7.

23. Hicks L, van der Graaf CM, Childress J, Cook E, Schmidt K, Rosenzweig F, et al. Streamlined preparation of genomic DNA in agarose plugs for pulsed-field gel electrophoresis. Journal of Biological Methods [Internet]. 2018 Mar 9; Available from: http://www.jbmethods.org/jbm/article/view/218/198

24. Coyle S, Kroll E. Starvation induces genomic rearrangements and starvation-resilient phenotypes in yeast. Mol Biol Evol. 2008 Feb;25(2):310–8.

25. García V, Rivera J, Contreras A, Ganga MA, Martínez C. Development and characterization of hybrids from native wine yeasts. Braz J Microbiol. 2012 Apr;43(2):482–9.

26. Struhl K. Direct selection for gene replacement events in yeast. Gene. 1983 Dec;26(2–3):231– 41.

27. Huxley C, Green ED, Dunham I. Rapid assessment of S. cerevisiae mating type by PCR. Trends Genet. 1990 Aug;6(8):236.

28. Coluccio A, Neiman AM. Interspore bridges: a new feature of the Saccharomyces cerevisiae spore wall. Microbiology (Reading). 2004 Oct;150(Pt 10):3189–96.

29. Lysis of viable yeast cells by enzymes of Arthrobacter luteus - PubMed [Internet]. [cited 2025 July 31]. Available from: https://pubmed.ncbi.nlm.nih.gov/5166255/

30. Strober W. Trypan Blue Exclusion Test of Cell Viability. Curr Protoc Immunol. 2015 Nov 2;111:A3.B.1-A3.B.3.

31. Ramírez F, Ryan DP, Grüning B, Bhardwaj V, Kilpert F, Richter AS, et al. deepTools2: a next generation web server for deep-sequencing data analysis. Nucleic Acids Res. 2016 July 8;44(W1):W160–165.

32. Afgan E, Baker D, Batut B, van den Beek M, Bouvier D, Cech M, et al. The Galaxy platform for accessible, reproducible and collaborative biomedical analyses: 2018 update. Nucleic Acids Res. 2018 July 2;46(W1):W537–44.

33. Robinson JT, Thorvaldsdóttir H, Winckler W, Guttman M, Lander ES, Getz G, et al. Integrative genomics viewer. Nature Biotechnology. 2011;

34. Jensen R, Sprague GF, Herskowitz I. Regulation of yeast mating-type interconversion: feedback control of HO gene expression by the mating-type locus. Proc Natl Acad Sci U S A. 1983 May;80(10):3035–9.

35. Soares EV. Flocculation in Saccharomyces cerevisiae: A review. Journal of Applied Microbiology. 2011;110(1):1–18.

36. Weiss EL. Mitotic exit and separation of mother and daughter cells. Genetics. 2012 Dec;192(4):1165–202.

37. Olson BJSC. Multicellularity: From brief encounters to lifelong unions. Elife. 2013 Dec 24;2:e01893.

38. Lander ES, Linton LM, Birren B, Nusbaum C, Zody MC, Baldwin J, et al. Initial sequencing and analysis of the human genome. Nature. 2001 Feb 15;409(6822):860–921.

39. Biggins S, Bhalla N, Chang A, Smith DL, Murray AW. Genes involved in sister chromatid separation and segregation in the budding yeast Saccharomyces cerevisiae. Genetics. 2001 Oct;159(2):453–70.

40. Brachmann CB, Davies A, Cost GJ, Caputo E, Li J, Hieter P, et al. Designer deletion strains derived from Saccharomyces cerevisiae S288C: a useful set of strains and plasmids for PCR-mediated gene disruption and other applications. Yeast. 1998;14(2):115–32.

41. Hammerschmidt K, Rose CJ, Kerr B, Rainey PB. Life cycles, fitness decoupling and the evolution of multicellularity. Nature. 2014 Nov 6;515(7525):75–9.

42. Fisher RA. The Genetical Theory of Natural Selection. Oxford, U.K.; 1930. 272 p.

43. Hammerschmidt K, Landan G, Domingues Kümmel Tria F, Alcorta J, Dagan T. The Order of Trait Emergence in the Evolution of Cyanobacterial Multicellularity. Genome Biol Evol. 2021 Feb 3;13(2):evaa249.

44. Bingham EP, Ratcliff WC. A nonadaptive explanation for macroevolutionary patterns in the evolution of complex multicellularity. Proceedings of the National Academy of Sciences. 2024 Feb 13;121(7):e2319840121.

45. Lyons NA, Kolter R. On the evolution of bacterial multicellularity. Current Opinion in Microbiology. 2015;24:21–8.

46. Brunet T, King N. The Origin of Animal Multicellularity and Cell Differentiation. Dev Cell. 2017 Oct 23;43(2):124–40.

47. Lindsey CR, Rosenzweig F, Herron MD. Phylotranscriptomics points to multiple independent origins of multicellularity and cellular differentiation in the volvocine algae. BMC Biol. 2021 Aug 31;19(1):182.

48. Feng B, Lin Y, Zhou L, Guo Y, Friedman R, Xia R, et al. Reconstructing Yeasts Phylogenies and Ancestors from Whole Genome Data. Sci Rep. 2017 Nov 9;7(1):15209.

49. Ratcliff WC, Herron MD, Howell K, Pentz JT, Rosenzweig F, Travisano M. Experimental evolution of an alternating uni- and multicellular life cycle in Chlamydomonas reinhardtii. Nature Communications. 2013;4.

50. Wang Y, Shirogane T, Liu D, Harper JW, Elledge SJ. Exit from exit: resetting the cell cycle through Amn1 inhibition of G protein signaling. Cell. 2003 Mar 7;112(5):697–709.

51. Oud B, Guadalupe-Medina V, Nijkamp JF, de Ridder D, Pronk JT, van Maris AJA, et al. Genome duplication and mutations in ACE2 cause multicellular, fast-sedimenting phenotypes in evolved Saccharomyces cerevisiae. Proceedings of the National Academy of Sciences. 2013 Nov 5;110(45):E4223–31.

52. 52. https://www.ncyc.co.uk/catalogue/sgrp-strain-set-2.

53. Doulcier G, Remigi P, Rexin D, Rainey PB. Evolutionary dynamics of nascent multicellular lineages. Proc Biol Sci. 292(2045):20241195.

54. Szyszka-Mroz B, Ivanov AG, Trick CG, Hüner NPA. Palmelloid formation in the Antarctic psychrophile, Chlamydomonas priscuii, is photoprotective. Front Plant Sci. 2022;13:911035.

55. Suwannachuen N, Leetanasaksakul K, Roytrakul S, Phaonakrop N, Thaisakun S, Roongsattham P, et al. Palmelloid Formation and Cell Aggregation Are Essential Mechanisms for High Light Tolerance in a Natural Strain of Chlamydomonas reinhardtii. Int J Mol Sci. 2023 May 6;24(9):8374.

56. Chen ICK, Khatri S, Herron MD, Rosenzweig F. Genetic predisposition towards multicellularity in Chlamydomonas reinhardtii. Genome Biol Evol. 2025 May 17;evaf090.

57. Kapsetaki SE, Cisneros LH, Maley CC. Cell-in-cell phenomena across the tree of life. Sci Rep. 2024 Mar 29;14(1):7535.

58. Meussdoerffer F, Tortora P, Holzer H. Purification and properties of proteinase A from yeast. Journal of Biological Chemistry. 1980 Dec 25;255(24):12087–93.

59. Hayashi R, Moore S, Stein WH. Serine at the Active Center of Yeast Carboxypeptidase. Journal of Biological Chemistry. 1973 Dec;248(24):8366–9.

60. Smeekens JM, Xiao H, Wu R. Global Analysis of Secreted Proteins and Glycoproteins in Saccharomyces cerevisiae. J Proteome Res. 2017 Feb 3;16(2):1039–49.

61. Pugh TA, Shah JC, Magee PT, Clancy MJ. Characterization and localization of the sporulation glucoamylase of *Saccharomyces cerevisiae*. Biochimica et Biophysica Acta (BBA) - Protein Structure and Molecular Enzymology. 1989 Feb 23;994(3):200–9.

62. Merkel O, Fido M, Mayr JA, Prüger H, Raab F, Zandonella G, et al. Characterization and function in vivo of two novel phospholipases B/lysophospholipases from Saccharomyces cerevisiae. J Biol Chem. 1999 Oct 1;274(40):28121–7.

63. Gancedo JM. Control of pseudohyphae formation in Saccharomyces cerevisiae. FEMS Microbiol Rev. 2001 Jan;25(1):107–23.

64. Martínez C, Cosgaya P, Vásquez C, Gac S, Ganga A. High degree of correlation between molecular polymorphism and geographic origin of wine yeast strains. Journal of Applied Microbiology. 2007 Dec 1;103(6):2185–95.

65. Jauert PA, Jensen LE, Kirkpatrick DT. A novel yeast genomic DNA library on a geneticin-resistance vector. Yeast. 2005;22(8):653–7.

66. Voth WP, Olsen AE, Sbia M, Freedman KH, Stillman DJ. ACE2, CBK1, and BUD4 in Budding and Cell Separation. Eukaryotic Cell. 2005;

67. Kuranda MJ, Robbins PW. Cloning and heterologous expression of glycosidase genes from Saccharomyces cerevisiae (a-mannosidase/glucanase/chitinase/Schizosaccharomyces pombe). Biochemistry. 1987;

68. MacCallum DM, Findon H, Kenny CC, Butler G, Haynes K, Odds FC. Different consequences of ACE2 and SWI5 gene disruptions for virulence of pathogenic and nonpathogenic yeasts. Infection and Immunity. 2006;

